# EEG Decoding Reveals the Temporal Dynamics and Functional Relevance of Goal-Relevant Representations

**DOI:** 10.1101/219741

**Authors:** Jason Hubbard, Atsushi Kikumoto, Ulrich Mayr

## Abstract

Models of action control assume that abstract task-set settings regulate lower-level stimulus/response representations. Yet, we know little about the functional and dynamic properties of task-set representations in humans. Using a cued task-switching paradigm, we show that information about task sets and lower-level stimulus/response aspects can be extracted through decoding analyses from the scalp electrophysiological signal (EEG) on the single-trial level and with high temporal resolution. Task-sets are active throughout the entire processing cascade and trial-to-trial variations in task-set strength emerges as a remarkably strong predictor of variability in performance, both within and between individuals. Also, taskset strength is related to stimulus representation strength at an early period and to the strength of response representations at a later period, consistent with the notion that task-sets coordinate successive, lower-level representations in a concurrent manner. These results demonstrate a powerful approach towards uncovering stages of information processing and their relative importance for performance.

---

The efficiency of information processing differs both moment to moment, and from one individual to the next. Such variability could reflect the quality of low-level, stimulus or response representations. Alternatively, it may arise from the strength of abstract, task-set representations that instantiate or control lower-level processes (1-3). For example, in the experimental paradigm we used in the current work (see Figure 1a), participants were informed on each trial through auditory cues, which of two tasks to perform (4, 5). For the color task, they attended to the color singleton within the array of objects and responded via button press whether the exact color was orange or purple. Similarly, for the orientation task, participants attended the orientation singleton and responded whether the line tilted to the left or to the right. In this situation, successful performance requires lower-level representations of the task cue, of the target location, and of the task-relevant feature/response. However, it may also require abstract task-set representations that differentiate between the color task context and the orientation task context and that ensure an adequate configuration of lower-level representations.

**Figure 1.**
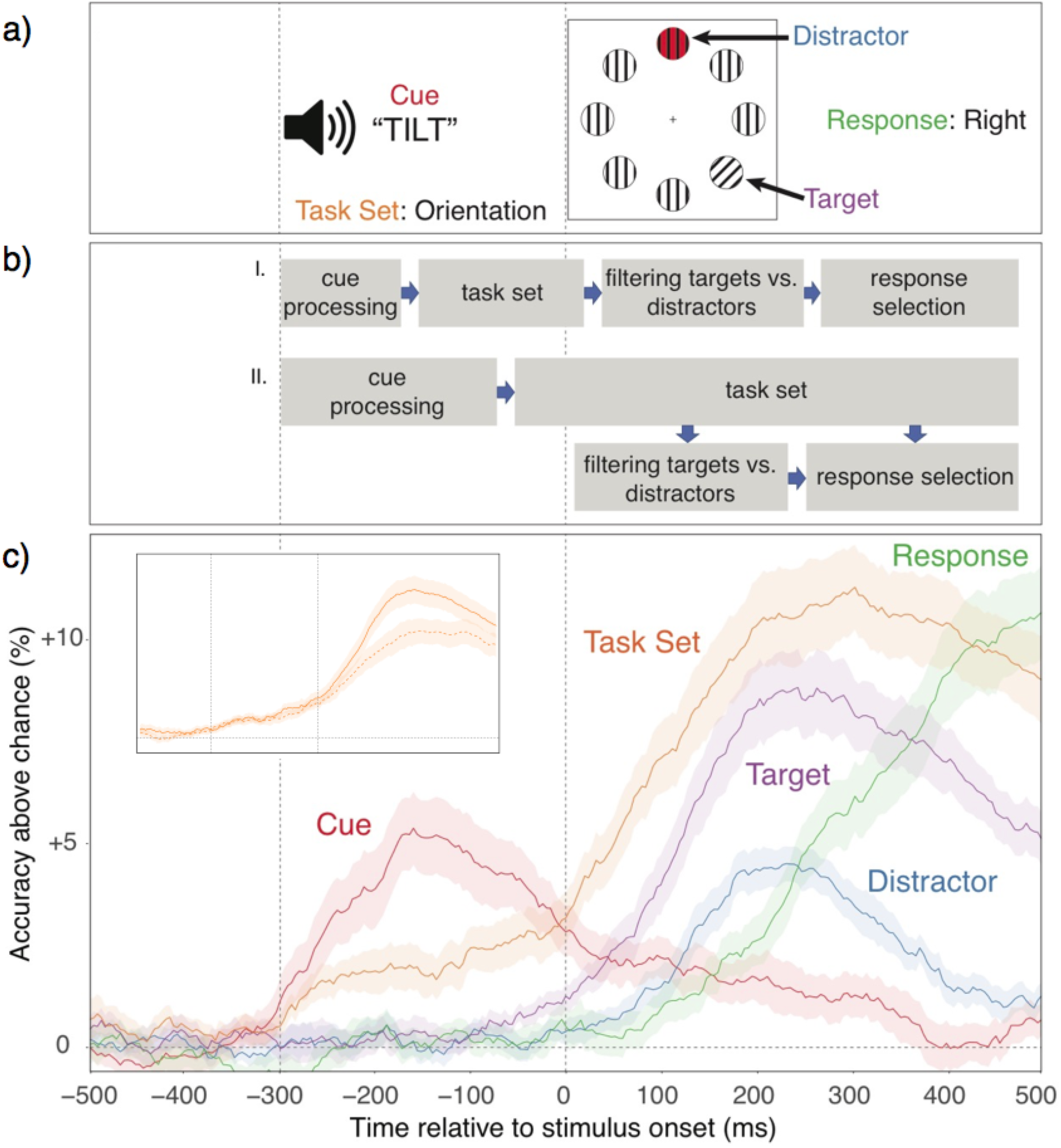
a) Stimulus timeline with each relevant task aspect. b) Competing models specifying either a (I) “preconfiguration” or a (II) parallel-activation relationship between task set and stimulus/response representations. c) Decoding accuracy of each aspect across time, relative to chance (*p*=.5, except for target and distractor, where it was *p*=.25). In all figures, shaded regions specify 95% within-subject confidence intervals. The insert shows how task-set decoding accuracy generalizes both within (filled line) and across target locations/responses (dotted line). Note, that in the current work we are particularly interested in within-individual variability in decoding accuracy and therefore the values presented here are based on averaged, trial-by-trial results. When performing decoding analyses based on averaged data, much higher decoding accuracy (>80% for some aspects) can be achieved.

Even though the existence of higher-level, control representations is a common assumption in models of cognitive control (3, 6-8) there are many open questions about the degree to which abstract task sets regulate performance at all, and how such regulation is achieved (4, 5, 9, 10). For example, as an alternative to the view that abstract, rule-like representations are necessary to modulate lower-level processes, some authors have pointed out that when unambiguous, environmental stimuli (i.e., cues) distinguish between competing response options, superficial cue representations could be sufficient to constrain lower-level processes (4). It is also currently not clear when exactly higher-level control occurs (see Figure 1b). Cue or task-set representations might be necessary to set up and preconfigure lower-level representations (11, 12). Alternatively, task sets may also become relevant only as competition between lower-level representations arises, in order to mold these representations in a goal-appropriate manner (3) (i.e., see the Parallel Activation account in Figure 1b). Addressing these and related issues requires methods that directly probe the status of goal-relevant representations with high temporal resolution. In particular, it would be important to examine not only when a specific representation is active, but to what degree it actually impacts performance.

Existing approaches, such as chronometric analyses of response-time (RT) patterns (13), the analysis of averaged evoked EEG (14), or fMRI BOLD signals (15) are of limited value for capturing temporal dynamics, or trial-to-trial variability, in the strength of different task-related representations. Moreover, it is of particular theoretical importance to characterize the dynamic behavior of abstract task-set representations. Yet, because such representations are not tied to specific stimuli or responses, they are difficult to pin down with the existing methods.

Recent work has suggested that a surprising amount of information about currently active representations can be extracted from EEG signals (16-19). Therefore, we applied a bottom-up, multivariate decoding approach to EEG data from the task-switching paradigm presented in Figure 1a. We extracted from the EEG signal information about each of the five, potentially relevant aspects (superficial cues, target and distractor locations, target feature/response, and task set). The decoding accuracy for each aspect provided highresolution information about the temporal dynamics and the relative importance of both lower-level, stimulus/response, and higher-level task-set representations.

## Results

### Representational Dynamics

Figure 1c shows that decoding accuracy unfolds consistent with standard expectations about the flow of information--from cue encoding, to task-set activation, to relevant and irrelevant stimulus locations, and finally to response codes. Remarkably, task-level information is decodable with high accuracy throughout almost the entire duration of the trial.

As mentioned, one prominent model suggests that the cognitive system does not actually rely on abstract task settings, but—at least when available—uses superficial cue representations to resolve ambiguity between competing, stimulus/response representations (4, 9). However, we find that cue decoding accuracy (i.e., discriminating between the two cues for each task) peaks during the pre-stimulus phase, but declines sharply once the stimulus is presented. This result is consistent with the view that cue representations are used to activate task set representations (5, 20), and are less involved with actually regulating task-specific processes.

But how exactly are tasks instantiated? It task-set representations are critical for “preconfiguring” lower-level processes, stimulus representations would need to wait until the task-set is firmly established (11, 12). Alternatively, task sets may be activated in parallel to low-level stimulus/response selection processes, biasing these in a goal-relevant direction (3). As shown in Figure 1c, there is limited task decoding during the prestimulus phase, but a substantial increase emerges once stimulus/response information becomes decodable. Thus, while the presence of prestimulus task-set information indicates some role for preconfiguration, the overall pattern suggests that—consistent with the parallel-activation idea—task-set representations become particularly critical once competition between conflicting stimulus/response representations needs to be resolved (see Figure 1b).

An important question to address is to what degree task-set decoding accuracy actually reflects abstract representations, rather than task differences in lower-level stimulus or response-selection processes. In fact, as reported in Figures S2 and S3, there are substantial task-dependent differences in lower-level features, which are probably tied to the differential saliency of color versus orientation stimuli. In principal, such lower-level differences could account for the task-decoding results. However, when we probed to what degree task-set decoding generalizes across target locations and features/responses (see SM), we found reliable, though somewhat reduced generalization of task-set representations (insert in Figure 1c). These results confirm that at least a substantial part of the decodable task-related information is indeed of a relatively abstract nature.

### Determining the Relevance of Representations

Going beyond average activation trajectories, the trial-by-trial decoding approach allows us to determine at what point in the trial, which of the represented aspects drive performance. To this end, we entered trial-by-trial classifier confidence associated with each of the five task aspects as simultaneous predictors of trial-by-trial RT variability within subjects, using mixed regression models for each timepoint (see SI for details). The coefficients shown in Figure 2a represent the unique predictive power associated with each aspect, as a function of time in the trial. Consistent with the view that task sets, and not superficial cue representations, control lower-level representations, we find that cue-related activity is largely irrelevant for performance, a result that also holds up when cue decoding accuracy is entered as the sole predictor. In contrast, task-set information becomes highly predictive of RTs during the post-stimulus phase, suggesting that fluctuations in the quality of task-set representations are a major source of trial-to-trial variability in performance. Further, the fact that the task-sets begin to predict RTs only once also the (independent) predictive power of stimulus and response information emerges, is consistent with the parallel-activation account (see also Figure S4).

**Figure 2.**
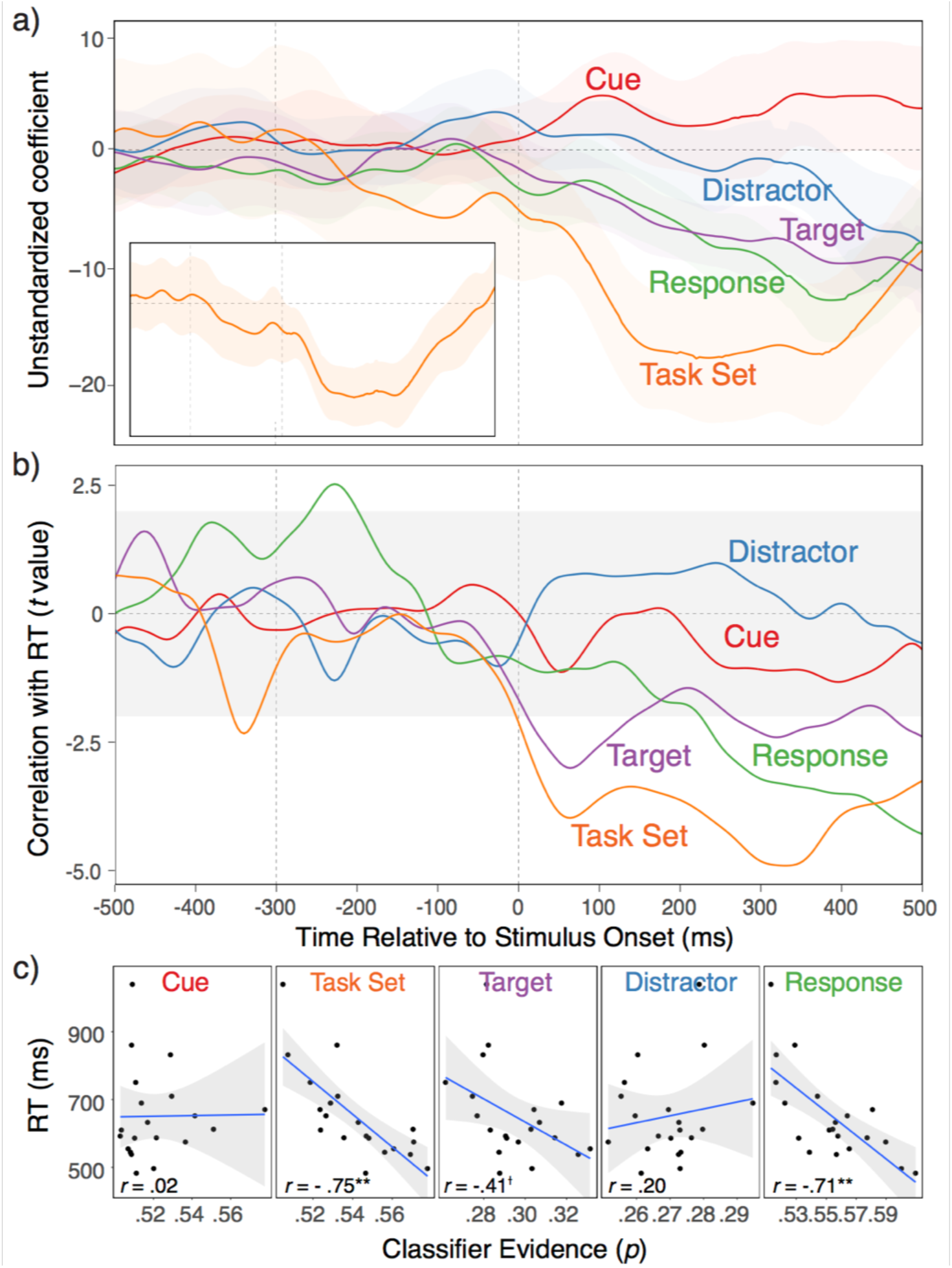
Within and between-individual relationships between decodability of all task aspects and RTs. **a)** Coefficients from multilevel, linear models with logit-transformed classifier evidence from all task aspects for a given timepoint simultaneously predicting RTs. The insert shows the coefficient when the task-set predictor is based on the generalization scores presented in the insert to Figure 1c. **b)** T-values representing simple correlations between individuals’ average, logit-transformed classifier evidence for each task aspect and their average RT. **c)** Scatterplots of relationships between subject-averaged classifier evidence and RTs during peak, average decoding accuracy periods for each task aspect (see Supplemental Material for details). For task-set generalization scores (see insert to Figure 1c), the correlation remained very robust at .63 (*p*<.01).

Again, we need to ask to what degree the predictive power of task sets is associated with abstract task-set representations rather than with lower-level differences between tasks. Therefore, we used instead of task-set classifier confidence the degree of generalization (see also insert to Figure 1c) to predict RTs. Indeed, as shown in the insert to Figure 2a, the generalizable aspect of the decoded task-set representation remains a robust predictor of performance.

We can also look at the degree to which the decoding accuracy for the different aspects is related to individual differences in RTs. In fact, we found that the temporal pattern of simple correlations between individuals’ average decoding accuracy for each task aspect and their average RT is very similar to the within-individual predictive pattern. As shown in Figure 2b, decoding accuracy for task sets is a major source of individual differences in RTs for nearly the entire post-stimulus period, whereas stimulus location correlates early and the feature/response aspect late in the post-stimulus phase (Figure 2b). In addition, Figure 2c shows the scatterplots for the correlations at each task aspects’ peak decoding accuracy (see Figure 1c). The current experiment was not designed to examine individual differences. However, confidence in these results is strengthened by the fact that the relationships are strikingly robust and highly consistent with the within-individual relationships.

### Relationship Dynamics between Representations

The general notion that cues activate task sets, which in turn bias stimulus and eventually response representations, leads to a straightforward prediction about the sequence in which different representations should be related to each other. To test these predictions we can examine the degree to which the decoding accuracy for different representations is coupled as a function of time in the trial. Figure 3 presents for each timepoint the relationships between the classifier confidence for task sets and each of the other aspects (to avoid clutter, we omit the distractor here, for which the relationship was close to zero throughout). As expected, early in the prestimulus phase, the strength of task representation is coupled with the strength of cue representation, likely indicating the retrieval of the task set based on the cue (5). Following stimulus onset, a correlation with the target location emerges and subsequently, a correlation with the response information. This pattern is again consistent with the parallel-activation account, where task sets coordinate lower-level representations in a concurrent manner (see Figure 1b).

**Figure 3.**
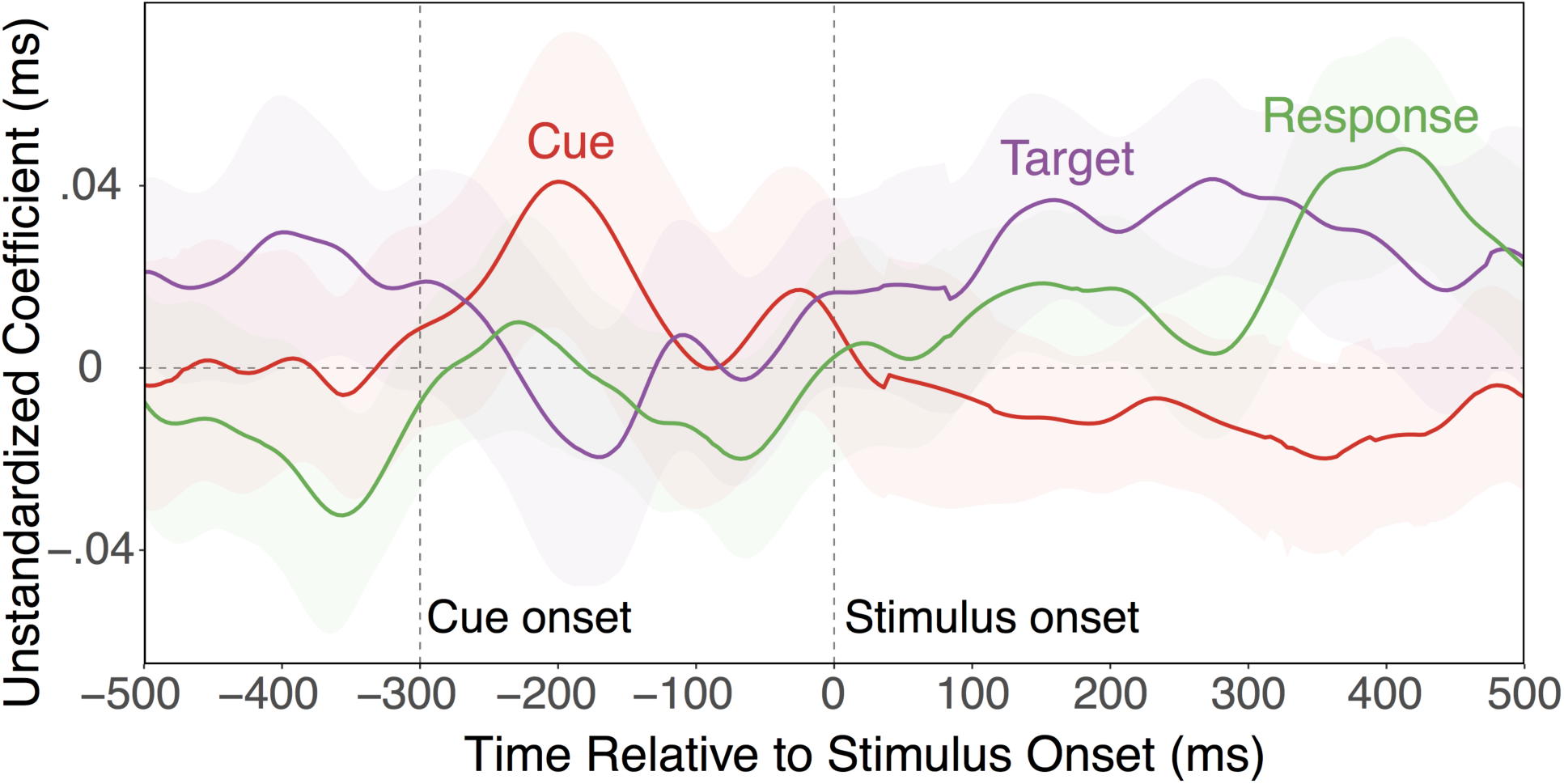
Coefficients representing the independent relationships between classifier confidence for task set on the one hand, and for cue, target, and responses on the other. For clarity of presentation, the relationship with distractor classifier confidence was omitted here, which hovered around 0 throughout.

### Effects of Task Switching

In the results presented so far, we had ignored potential effects of trial-to-trial changes in tasks. In fact, our version of the task-switching paradigm was optimized towards EEG decoding analyses, not towards producing large switch effects (i.e., both the use of long inter-trial intervals and of spatially separate, task-related features can be expected to reduce between-task competition). Indeed, behavioral switch costs were small (for details see Supplemental Material). Nevertheless, we examined the degree to which the switch/repeat contrast plays out in the decoding results. We constrained our analyses a-priori to the 150 ms intervals centered around the peak of the activation trajectories for each feature (see Figure 1c). In addition, given the strong relationship between RTs and decoding accuracy for task, stimulus locations, and response, we also conducted a median split into fast and slow RT trials. The median-split was conducted within each subject, task, and switch condition; values were than averaged across tasks and subjects, but presented separately for no-switch and switch trials. The dominant aspect in Figure 4 is again the strong relationship between RTs on the one hand and task, target location, and response representations on the other. In addition, switching tasks leads to weakened task-set representations, both in general (switch main effect: F[1,19]=6.69, p=.018), but in particular on slow-RT trials (fast/slow x switch interaction: F[1,19]=6.62, p=.019). Also, distractor representations were increased on switch trials, F(1,19)=5.23, p=.034). Thus, at the time of peak task-set activation, the task-related information is less robust on switch than on no-switch trials, whereas information related to the competing task is more strongly expressed.

**Figure 4.**
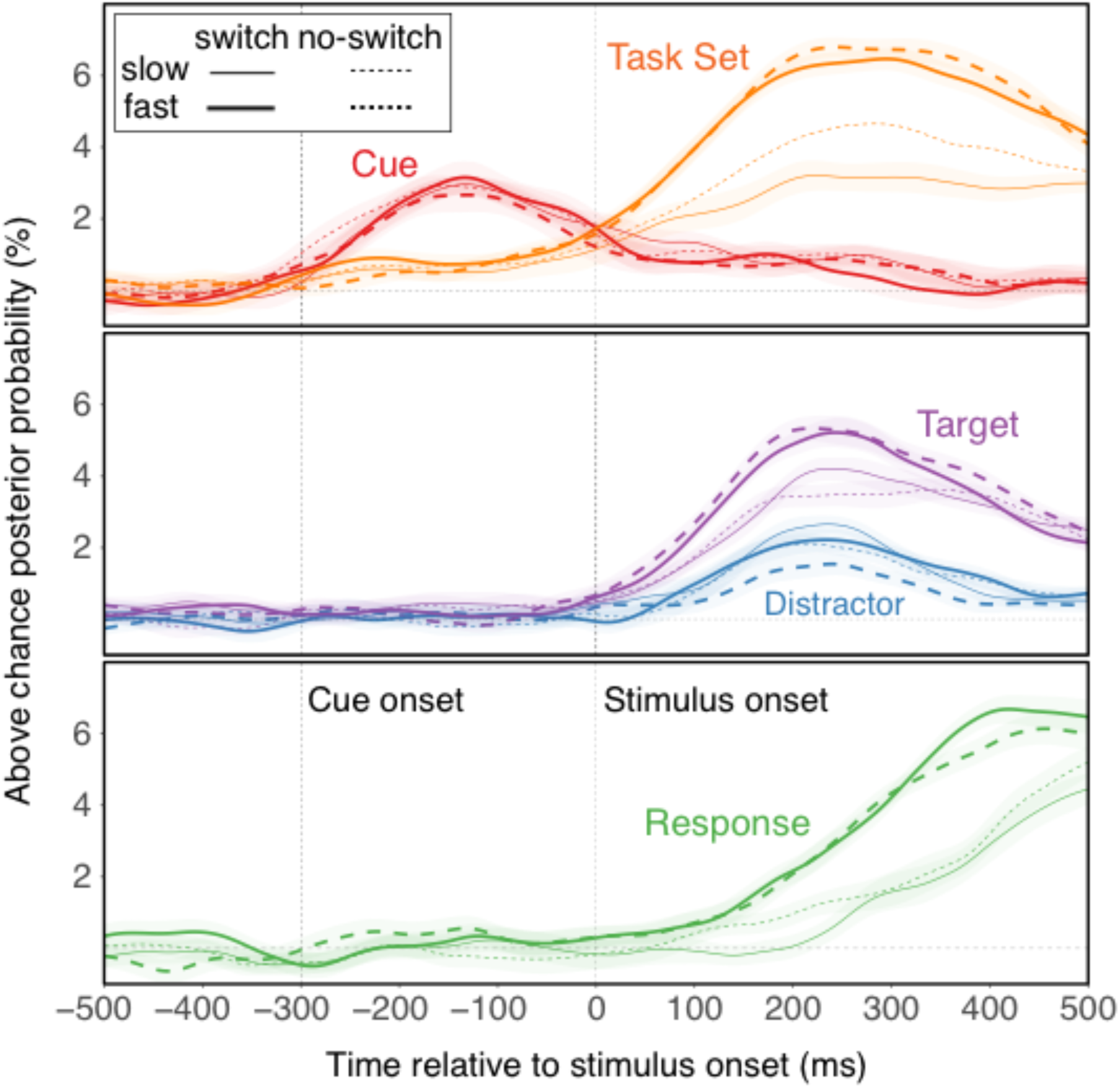
Average classifier confidence, separately for switch vs. no-switch trials and fast vs. slow RTs (determined via median split within individuals and switch vs. no-switch trials).

## Discussion

When people need to respond to a given stimulus in a flexible, context-dependent manner, the flow of information processing cannot rely on sensory or response representations alone. Rather, stimulus and response selection needs to be constrained by representations of the environmental context (i.e., cues) and potentially of the current stimulus-response rules (i.e., task sets). There is a substantial behavioral (1) and fMRI neuroimaging (21, 22) literature on how we select and change such higher-level representations in humans. However, these methods have not provided a precise account of the place and the relevance of cue or task-set representations within the overall information processing cascade.

We made progress on these questions by decoding information about key goal-relevant representations from EEG signals. Our results reveal a plausible sequence of active representations of target and distractor locations, as well as of features or response choices. For example, the timing of the target/distractor location representations is highly consistent with recent work using eye-tracking to assess the dynamics of attentional allocation to task-relevant and irrelevant features (12, 23). More importantly, our results also reveal the time course for both cue and task representations. Task cues were highly decodable as soon as the cue was presented during the prestimulus phase, but were less strongly expressed once the stimulus appeared. In contrast, task representations exhibited the reverse pattern, with limited activity during the post-cue/pre-stimulus phase, but a very strong presence during the entire stimulus-to-response. This result is consistent with findings from single-unit or multi-unit recoding work from primates, which has suggested that representations of the larger behavioral context bridge across the entire, lower-level processing cascade (24, 25).

The pattern of average, cue and task activation trajectories provides initial, theoretically relevant information. For example, the fact that task-set decoding accuracy is much higher than cue decoding accuracy (at least after stimulus presentation) suggests that--contrary to one prominent model (4)--task-set activity is more important than superficial cue information in controlling lower-level representations. Yet, average decoding accuracy allows no from conclusions about the functional relevance of the different representations. This is where decoding scores on the single-trial level yield important, additional information. Specifically, we used these scores to predict trial-to-trial variability in RTs, and thereby determined in a time-resolved manner, which representations drive performance (Figure 2). Interestingly, the pattern of predictive relationships indicates that cue representations do not explain variability in performance. In contrast, task representations emerge as a very strong predictor of RTs and explain substantial variability both within and across individuals. The explanatory power of task representations is again largely limited to the post-stimulus phase and is most robust when also stimulus and response effects are particularly strong. Interestingly, task-set information (but not cue information) also emerged as a very strong predictor of inter-individual differences in performance. Moreover, an analysis of interrelationships between task-set and lower-level representations (see Figure 3) indicate that task sets were coupled in the post-stimulus phase initially with the target-location representations, and thereafter also with feature/response representations. Combined, these results strongly suggest that lower-level representations are configured through relatively abstract task or attentional settings, not through superficial cue representations. Further, the trajectory of average representational strength and the predictive pattern is most consistent with the parallel activation model (Figure 1b), where task sets can shape lower-level processes in a concurrent manner (3).

Our results cannot be taken to rule out the possibility of functionally relevant, preparatory activity before stimulus/response processing sets in. In fact, the use of cue information to retrieve the current task set is a necessary process that clearly happens within the cue-to-stimulus interval (5). The early inter-relationship between strength of cue and task representations likely is an expression of this retrieval activity (Figure 3). It is reasonable to assume that the 300 ms interval between cue and stimulus interval was sufficient to absorb major, within-individual or between-individual variability in the duration or quality of this process, thus preventing any predictive effects of cue or task-set representations from revealing themselves. Also, it is very well possible that with longer preparatory intervals, greater proactive task-set activity may be observed. Both behaviorally and in fMRI neuroimaging studies, preparation effects are well documented (26-29). However, the fact that task-set representations were strongest, and also most predictive of performance in the presence of stimulus and response representations suggests that a key function of task sets is to regulate these lower-level representations in a concurrent manner. This conclusion is also consistent with a large body of behavioral work suggesting that task-selection costs cannot be easily eliminated through opportunity for preparation (1).

Several results indicate that the task decoding accuracy we observed does in fact represent the strength of relatively abstract, task or attentional settings. For example, in the predictive analyses, task-set classifier confidence explained substantial, within-individual variability in RTs over and above the predictive relationships between RTs and lower-level aspects. We also demonstrated that task representations generalized across target positions and features/responses. Furthermore, these generalization scores proved nearly as predictive of within-individual and between-individual variability in performance as the regular decoding accuracy. In the Supplemental Material we report an additional analysis, where we controlled in the predictive analyses (Figure 2) for the degree to which lower-level representations generalized across tasks. Again, we found no change to the overall predictive pattern, confirming that the decoded task-set representations were relatively abstract.

In a recent study by Siegel and colleagues (25), monkeys performed a task-switching paradigm while multi-unit activity was recorded in critical anatomical areas along the entire sensory-motor processing stream (see also, (30-32)). In several key aspects, the pattern of results was remarkably similar to the current findings. In particular, cue information was highly prominent during the prestimulus phase, but then tampered off in the post-stimulus phase. Task information emerged concurrently with cue information, but then increased dramatically as stimulus and response choice information was processed during the stimulus phase. The convergence of results across species and methods suggests that some of the same information that is conveyed through neural-level recordings can also be extracted through scalp EEG signals. The fact that we were able to extract information about task-relevant features through relatively sparse recordings from the scalp is generally consistent with the fact that in Siegel et al. both higher-level and lower-level aspects where represented throughout all cortical regions, albeit with varying strengths across regions(33, 34).

To summarize, ever since W. Wundt (35) and F.C. Donders (36), researchers have tried to characterize the cascade of goal-directed information processing and pinpoint the source of variability in processing efficiency in the human cognitive system. We show here that EEG-based, trial-by-trial decoding analyses can clarify the relative role and the temporal dynamics of both lower-level stimulus/response, as well as abstract task-set representations. In particular, the ability to pinpoint within the processing cascade the exact source of performance differences, is a unique feature of this approach. These and related methods help bridge the gap towards primate work that captures the information-processing flow in the activity of individual neurons.

## Methods

### Experimental Procedure

Twenty participants participated in this experiment. They were compensated at a rate of $10 per hour, with additional incentives based on performance on the task (see Supplemental Material for further details). All experimental procedures were approved through the University of Oregon’s Human Subject Review Board.

We used a cued task switching paradigm (27) that was closely modelled after a paradigm that we had previously used in the context of eye-tracking experiments ((12, 23); see Figure S1). On each trial, an auditory cue indicated which of the tasks, the Color task or the Orientation task, participants had to complete. Each task was paired with two auditory cues: “color” or “hue” for the Color task, and “tilt” or “lean” for the Orientation task. Following Monsell and Mizon (27), the two sets of cues (Set A=”color” and “tilt”, Set B=”hue” and “lean”) were alternated across consecutive trials. This ensured that task-switch costs are not contaminated by superficial cue-priming effects (4, 5) and that we could independently decode cue and task information(25).

The stimulus array consisted of 8 circular gratings (diameter of each ∽ 2.4 degrees) in a larger circular arrangement (diameter ∽ 12.5 degrees). The stimulus array always contained six, neutral stimuli consisting of vertical, black and white gratings. In addition, there was (a) one color singleton stimulus with a vertical grating shaded in one of two colors, either “yellowish” or “reddish” and one (b) orientation singleton with a black grating oriented either 30 degrees to the left or the right. For the Color task, participants had to attend to the color singleton and press the left (z) key for a “yellowish” target and the right (/) key for the “reddish” target. For the Orientation task, participants had to attend to the orientation singleton and press the left (z) key for a left-tilted target and the right (/) key for the right-tilted target.

Each trial began with a 700 ms prestimulus interval with a fixation cross in the center of the screen. The auditory cue was presented in the last 300 ms of this interval, so that the stimulus array appeared as soon as the auditory cue completed. Participants were instructed to respond as quickly and accurately as possible. The stimuli remained on the screen until a response was made. In case of a mistake, an error tone was emitted for 100 ms. During the following inter-trial interval (ITI), which was jittered between 750 and 937 ms, participants were instructed to blink before the next trial began.

The experiment began with two single-task practice blocks (one for each task, order counter-balanced), and a task-switching practice block (20 trials), followed by 22 test blocks of 64 trials each. All task aspects were determined randomly on a trial-by-trial basis. This includes the selection of tasks, yielding an average switch rate of *p*=.5.

Participants were seated approximately 70 cm from the screen, and instructed to keep their eyes at fixation and not blink throughout the trial; trials containing eye movements were detected via electrooculogram (EOG) and excluded from the analysis.

### EEG Decoding Analyses

After initial preprocessing and identification of artifacts (see SM), the single-trial EEG data were decomposed into a time-frequency representation via wavelet decomposition (for details see SM). For simplicity we focused on frequency bands that are most often presented in the literature: delta (2-3 Hz), theta (4-7 Hz), alpha (8-12 Hz), and beta (13-31 Hz). For each frequency band, we averaged the power signal across the range of interest. For each trial, we extracted a window centered around stimulus onset, starting 500 ms before and extending 500 ms after the onset. The end of this interval corresponds to the 70^th^ percentile of the RT distribution, ensuring that at least 70% of trials are still in progress at that point. As we were interested in the dynamics of the information in time, we performed the analyses separately for each 4 ms epoch (referred to as timepoint). Thus, for each timepoint and eachj of the 20 electrodes, the power in each of the four frequency bands, served as input for the decoding analyses (i.e., 80 features).

In the decoding analyses, we examined the extent to which the spatial pattern of the EEG power across the scalp and four frequency bands was predictive of each task aspect: cue (“color”/”hue” or “tilt”/“lean”, classified within tasks), task (Color or Orientation), target position (partitioned into 4 bins, coded 1-4), distractor position (bins 1-4), and the response (left vs. right). For the target/distractor location, we decoded positions based on the bin that each item appeared in (e.g., bin 1=top and top-right position, bin 2=right and bottom-right position, etc.). This ensured that the target and distractor occupied each combination of bins with equal frequency (including sharing identical bins), thus ensuring that successful distractor decoding is not simply due to the classifier decoding “not target position”. Note that in the current paradigm we cannot distinguish between the manual response and the specific target stimulus (e.g., left-tilted grating, or reddish grating) as they were confounded. As we wanted to isolate the discriminability of each aspect regardless of any task differences, we performed the decoding separately within each task (except, of course in decoding the task set itself). The results from these analyses were then averaged. Previous work has established that different types of information are encoded in brain oscillations at particular frequency bands, which motivated decomposing the raw EEG signal into the separate bands. However, in the present investigation we were agnostic to which bands encode which type of information, and thus concatenated all 4 bands together in the decoding analysis.

Prior to decoding, the EEG data were z-scored so that the mean of each trial’s data was 0 without baseline activity subtraction. We performed all analyses separately for each subject and then averaged results across subjects. The L2-regularized logistic regression was used, as implemented in the scikit-learn package in Python ((37)), with a tolerance of 1 x 10^-4^ and the inverse of the regularization strength (C) set to 1.0. Multi-class classification (which was the case for target and distractor positions), was implemented as a series of binary classifications. For all aspects, we used a 4-fold cross-validation procedure where 75% of trials were used in the training set, and the remaining 25% of trials were used as the test set, and this was repeated until each trial had an opportunity to be part of the test set.

For the main decoding analysis (Figure 1c), we reported the decoding accuracy, averaged within subject, time point, and factors of interest (e.g., task), and then across subjects. We used decoding accuracy here as it is most consistent with how such results are presented in the literature. In contrast, Figures 2-4 are based on classifier confidence, which provide a continuous prediction score on the trial-by-trial level.

## Acknowledgements

Jason Hubbard and Atsushi Kikumoto equally contributed to this work. This work was supported in part by National Institute of Aging grant R01 AG037564-01A1, NSF grant 1734264, as well by an Award by the Humboldt Foundation to Ulrich Mayr.

